# Urokinase receptor associates with TLR4 interactome to promote LPS response

**DOI:** 10.1101/2020.06.10.143826

**Authors:** Yulia Kiyan, Sergei Tkachuk, Anna Gorrasi, Pia Ragno, Inna Dumler, Hermann Haller, Nelli Shushakova

## Abstract

GPI-anchored uPAR is the receptor for the extracellular serine protease urokinase-type plasminogen activator (uPA). Binding of uPA to uPAR localizes proteolytic cascade activation at the cell surface and can induce intracellular signaling. As uPAR possesses no transmembrane domain, it relies on uPAR cross-talk with various membrane receptors. Though uPAR role in inflammatory processes is well documented, underlying mechanisms are not fully understood. In this study we demonstrate that uPAR is a part of Toll-like receptor 4 (TLR4) interactome. GPI-uPAR and soluble uPAR colocalized with TLR4 on the cell membrane and interacted with scavenger receptor CD36. We show that downregulation of uPAR expression resulted in diminished LPS-induced TLR4 signaling, less activation of NFκB, and decreased secretion of inflammatory mediators in myeloid and non-myeloid cells in vitro. In vivo uPAR−/− mice demonstrated strongly diminished inflammatory response and better organ functions in cecal ligation and puncture mouse polymicrobial sepsis model. Our data show that uPAR can interfere with innate immunity response via TLR4 and this mechanism represents a potentially important target in inflammation and sepsis therapy.

## Introduction

uPAR is the receptor for urokinase-type plasminogen activator (uPA), an extracellular serine protease and important activator of ubiquitous multifunctional protease plasmin. uPAR is anchored to the outer cell membrane leaflet via GPI anchor. Binding uPA to uPAR localizes proteolysis at the cell surface to facilitate spatially and temporally restricted activation of plasmin. Wide substrate specificity of plasmin provides for multiple functions of the protease such as fibrin cloth lysis, tissue remodeling, cell migration^1,2^. In addition, uPAR fulfills roles independent on the proteolytic activity of uPA. Binding of uPA or its catalytically inactive amino terminal fragment to uPAR or uPAR overexpression induces intracellular signaling pathways orchestrating important cellular functions such as proliferation, differentiation, migration, DNA repair^1,3,4^. Since uPAR is a GPI anchored protein and lacks a transmembrane domain, it relies on uPAR interaction with other receptors to transduce signals across cell membrane. uPAR interaction with several transmembrane receptors, integrins, and ECM components has been demonstrated^1^.

uPA/uPAR are expressed by many cells of hematopoietic origin^5^ and endothelial cells^6^,^7^. Expression of uPAR system can be rapidly upregulated in response to bacterial infection or inflammation. Despite the role of uPAR in inflammatory processes attracted attention^8,9^, its role is still not fully understood. Data obtained using uPAR−/− and uPA−/− mice models suggest that uPAR role in response to bacterial infection and innate immunity can be independent from uPA and its catalytic activity^10,11^. Effects of uPAR are often attributed to the deficient migration resulting in impaired infiltration of immune cells. Thus, uPAR−/− mice showed reduced accumulation of inflammatory cells in the lung upon *Streptococcus pneumoniae and Pseudomonas aeruginosa* infection^10,11^. This was accompanied by stronger propagation of the infection and higher mortality. Interestingly, *S. pneumoniae* caused modest increase in the lung levels of cytokines and chemokines in uPAR−/− mice. *S. pneumoniae* and its cell wall component lipoteichoic acid (LTA) are recognized primarily by TLR2 receptor^12^. In another study Liu and coworkers ^13^ addressed uPAR/TLR2 cross-talk directly. They reported that uPAR−/− neutrophils demonstrate diminished response to TLR2 ligand, PAM3CSK4 during in vitro stimulation. mRNA expression of cytokines in response to PAM3CSK4 was unchanged in uPAR−/− cells but the secretion of cytokines was decreased.

Sepsis is a severe and a life threatening condition that develops as an inadequate response to infection and may lead to organ dysfunction^14^. Plasma level of soluble uPAR (suPAR) alone and in combination with other biomarkers serves as a prognostic predictor and a marker in patients with sepsis and systemic inflammatory response (SIRS)^15,16^. We have previously demonstrated that uPAR cooperates with CD36 and TLR4 to mediate signaling induced by binding of oxidated low density lipoprotein (oxLDL) in vascular smooth muscle cells^17^. OxLDL is an important Danger Associated Molecular Pattern (DAMP) molecule regulating survival and phenotype of macrophages, endothelial cells, smooth muscle cells. We demonstrated that downregulation of uPAR in human vascular smooth muscle cells was protective against oxLDL-dependent phenotypic modulation. Scavenger receptor CD36 and innate immune receptor TLR4 also recognize Pathogen Associated Molecular Pattern molecules (PAMPs) such as LPS^18,19^ and play important roles in sepsis. We asked, if uPAR can interfere also with PAMPs signaling and whether interaction of uPAR with these receptors is important in vivo in sepsis.

## Results

### 1. uPAR−/− myeloid cells are less responsive to LPS stimulation

Whole blood collected from wild type (WT) and uPAR −/− mice was stimulated ex vivo with LPS from *E.coli*. After optimization of the stimulation conditions (Suppl. Fig. 1), the release of cytokines was assessed after 3 hrs of the blood stimulation with 50 ng/ml LPS using the Cytometric Bead Array and flow cytometry. As shown in Figure 1, LPS stimulation resulted in strongly increased levels of TNFα and IL-6 in plasma. However, this up-regulation was significantly decreased in the blood obtained from uPAR−/− in comparison to WT mice. The expression of IFNγ was also decreased in uPAR−/− blood though the difference has not reached the significance level (Suppl. Fig. 1B). The expression of MCP-1 and IL-12p70 were not increased by LPS and were similar in WT and uPAR−/− blood (Suppl. Fig. 1B). The expression of IL-10 on the contrary, was higher in uPAR−/− blood (Fig. 1A). Accordingly, IL6/IL-10 ratio was strongly decreased in uPAR−/− mice suggesting that knock-out blood cells demonstrated less inflammatory response.

**Figure 1.**
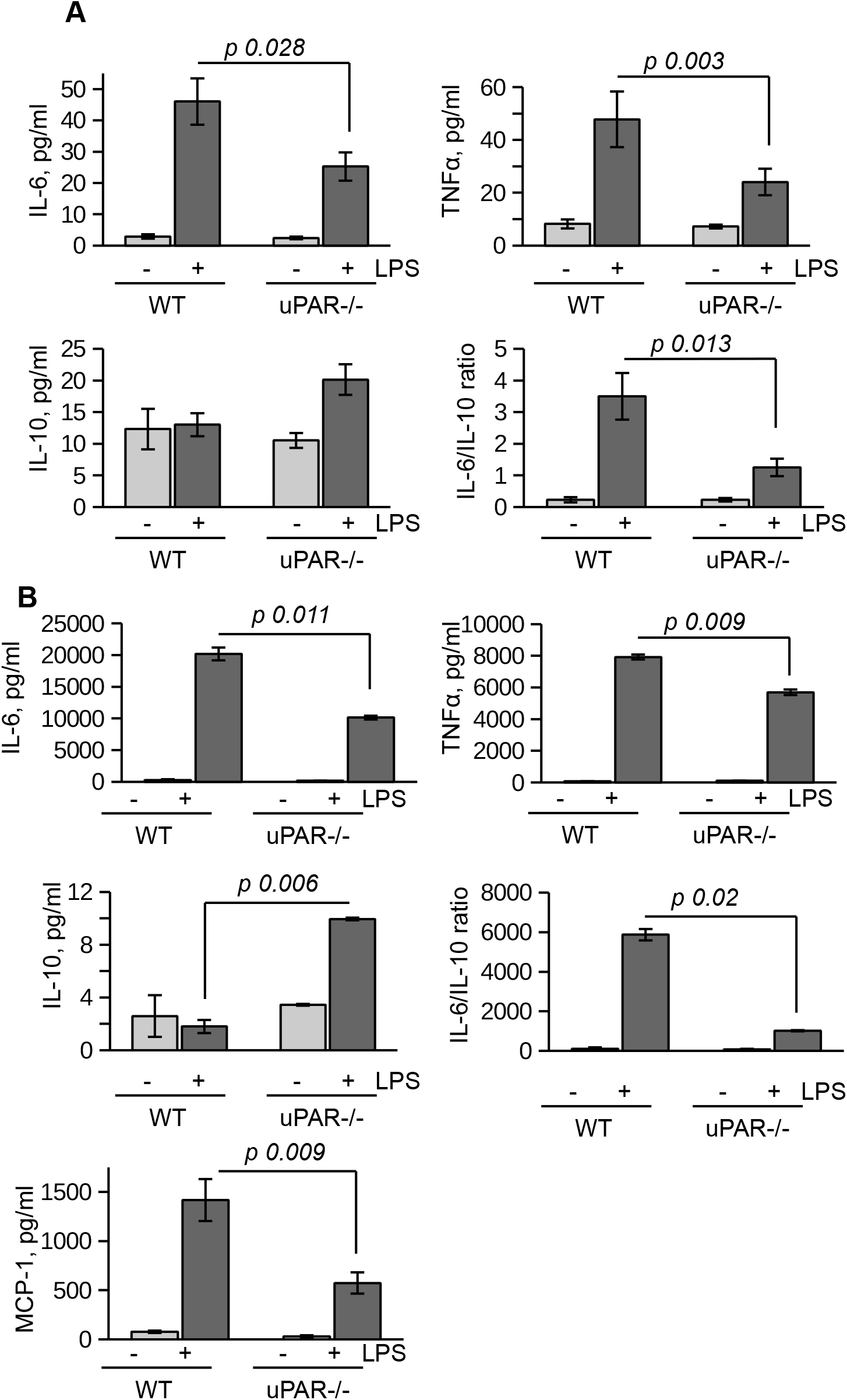
uPAR is essential for the response of myeloid cells to LPS. **A.** Whole blood from WT and uPAR−/− mice was stimulated ex vivo with 50 ng/ml LPS for 4 hrs at 37°C. Then, blood was centrifuged and cytokines content in plasma was assessed using mouse Cytometric Beads Array. **B.** Peritoneal macrophages isolated from WT and uPAR−/− mice were stimulated with 100 ng/ml LPS for 24 hrs. Expression of cytokines was assessed using beads array.

The whole blood response to LPS is initially mediated by the response of monocytic cells and largely mediated by TLR4^20^. Therefore, the model of the blood stimulation with LPS ex vivo implies uPAR participation in inflammatory signaling of TLR4 expressing monocytic cells. To investigate this opportunity further, we isolated primary peritoneal macrophages from uPAR−/− and WT mice. As shown in Figure 1B, uPAR−/− macrophages demonstrated significantly decreased expression of IL-6, TNFα, and MCP-1 after LPS stimulation compared to WT cells. Similar to whole blood stimulation, IL-6/IL-10 ratio was strongly decreased in uPAR−/− macrophages.

To assess the ability of uPAR to associate with proteins of TLR4 interactome, peritoneal macrophages from uPAR−/− mice were treated with mouse suPAR. Then the cells were fixed and stained for confocal microscopy. As shown in Figure 2A, in the presence of LPS suPAR also colocalized with membrane TLR4. Despite the ability to associate with TLR4 interactome, suPAR by itself had not induced any significant expression of IL-6 and TNFα in primary WT and uPAR−/− macrophages (Fig. 2B). However, suPAR promoted LPS response in uPAR−/− cells (Fig. 2B) confirming that suPAR can integrate into TLR4 interactome and this integration can have physiological relevance in LPS-induced response.

**Figure 2.**
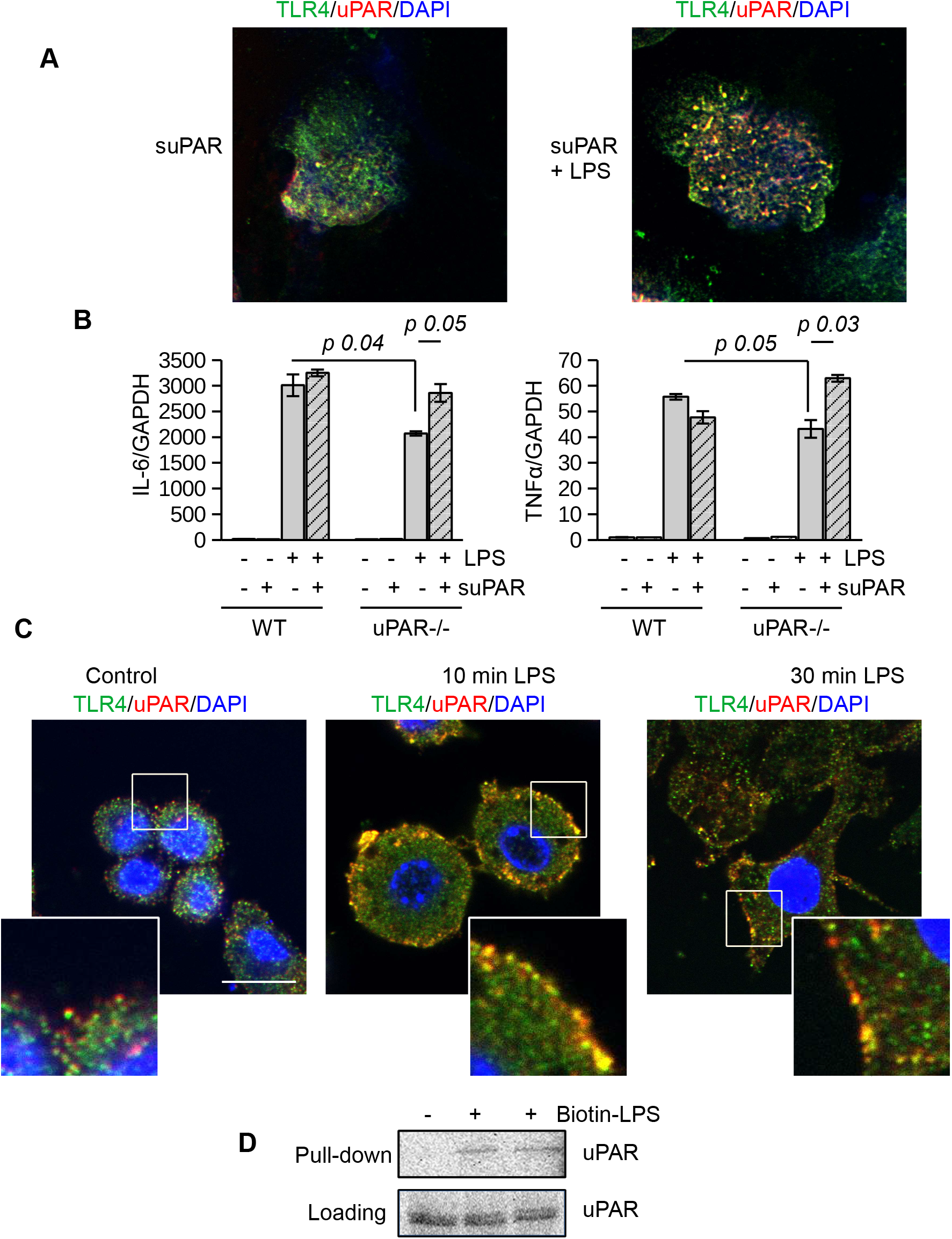
(s)uPAR is a part of TLR4 interactome. **A.** Primary peritoneal macrophages from uPAR−/− mice were stimulated with suPAR with or without LPS for 15 min. Then, cells were fixed and stained for TLR4 (Alexa 488, green) and uPAR (Alexa 647, red). DAPI was used as nuclear stain. Scale bar 10 μm. **B.** Primary WT and uPAR−/− macrophages were stimulated with 100 ng/ml LPS and1μg/ml suPAR for 3 hrs **C.** Raw 264.7 cells were stimulated with 100 ng/ml LPS, fixed and stained as in A. Scale bar 12.5 μm. **D.** Raw 264.7 were stimulated with 1μg/ml biotin-LPS for 30 min, then cell lysis was performed. Protein complexes were precipitated using Streptavidin magnetic beads and analyzed by western blotting using anti-murine-uPAR antibody.

Similar LPS-dependent co-localization of uPAR with TLR4 was observed using confocal microscopy in mouse Raw 264.7 macrophage cell line. These cells have been selected due to their high expression level of TLR4 and uPAR and demonstrate strong LPS response. As shown in Figure 2C, in unstimulated cells uPAR and TLR4 located in close proximity. In the presence of LPS, increased colocalization of the receptors was observed. This was further confirmed by biotin-LPS pull down assay performed in Raw 264.7 cells (Fig. 2D) - uPAR was found in the protein complex precipitated by biotin-LPS.

Together, this data showed that in the presence of LPS uPAR can integrate into signaling complex of TLR4 and promote the inflammatory response of myeloid cells.

### 2. LPS response of uPAR−/− non-myeloid cells is impaired

Non-myeloid cells also express TLR4 and its co-receptors and play important role in innate immunity response^21^. Mesothelial epithelium covers the internal body cavities and organs, and poses the first line of defense in abdominal bacterial sepsis. The expression of TLR4 by mouse mesothelial epithelial cells and their response to LPS have been reported^22^. In our experiments immortalized mouse mesothelial epithelial cells demonstrated a strong response to LPS by high upregulation of IL-6, TNFα, MIP-2, and MCP-1. We transfected mesothelial cells with control and murine uPAR siRNA (Fig. 3A) and assessed their response to LPS in vitro. As shown in Fig. 3B, the inflammatory response of uPARsi mesothelial cells was strongly impaired and the expression of IL-6, TNFα, CXCL2, and MCP-1 was decreased in comparison to the control cells.

**Figure 3.**
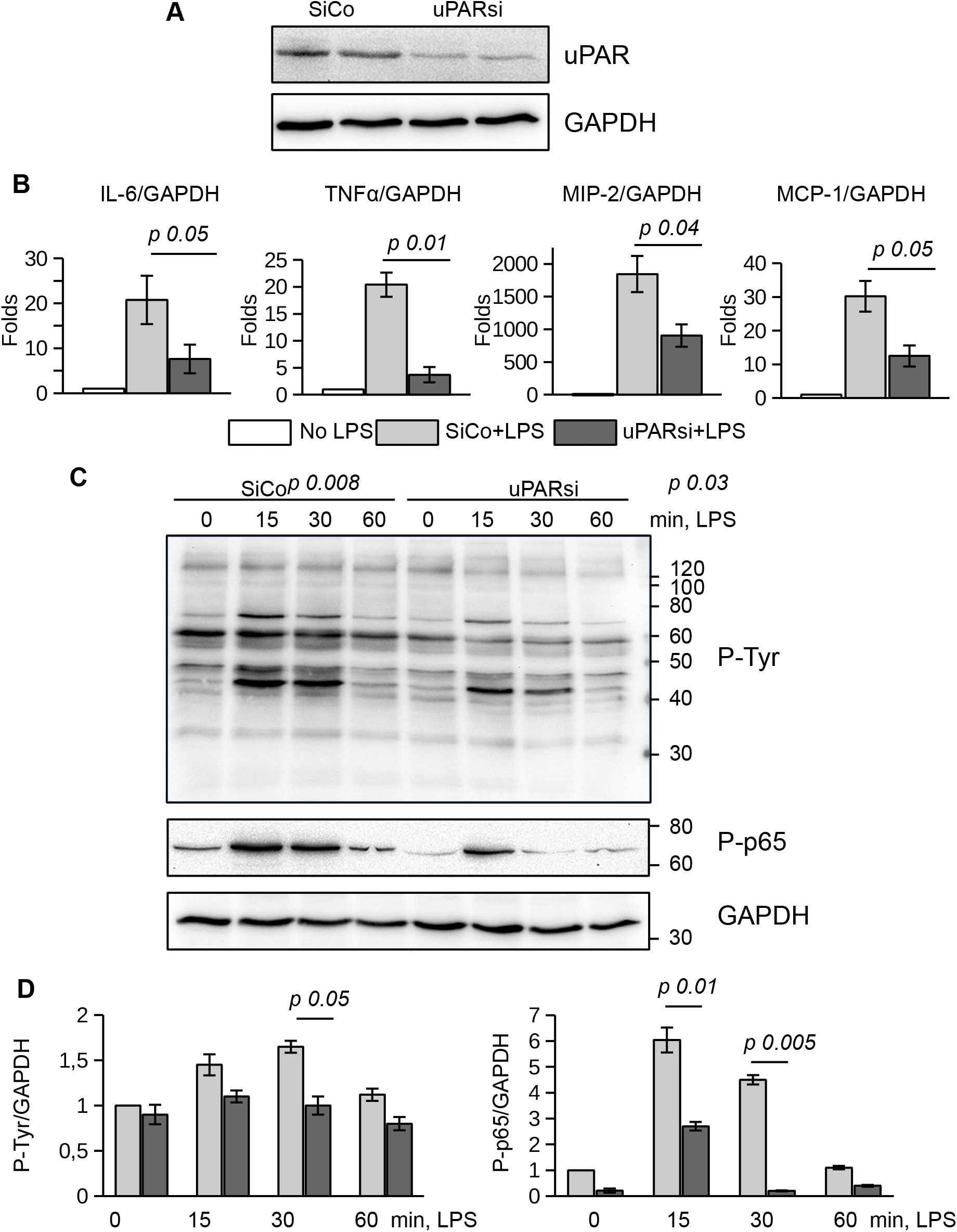
uPAR is essential for the response of mesothelial epithelial cells to LPS. **A.** Downregulation of uPAR expression in mouse mesothelial epithelial cells. **B.** LPS response of SiCo and uPARsi mouse mesothelial cells was assessed after stimulation with 100ng/ml LPS for 3 hrs. Expression was analysed by TaqMan RT-PCR. **C.** LPS-induced protein tyrosine phosphorylation was assessed in SiCo and uPARsi mouse mesothelial cells by western blotting of the whole cell lysate with P-Tyr antibody (upper panel) and P-p65 antibody (middle panel). GAPDH shows loading control (lower panel). **D.** Quantification of tyrosine (left) and p65 phosphorylation (right) from three independent western blotting experiments.

To assess mechanisms of uPAR interference with LPS-induced signaling, we investigated protein tyrosine phosphorylation in uPARsi mesothelial cells. As shown in Fig. 3C, D, tyrosine phosphorylation of multiple proteins was diminished in the absence of uPAR. In particular, phosphorylation of NFκB p65 was strongly decreased.

One of the most vulnerable organs affected by sepsis is the kidney. Recent data demonstrated that kidney tubular epithelial cells participate in immune response, express TLR4 and respond to LPS by expression of inflammatory cytokines^23^. To investigate the role of uPAR in the inflammatory response of kidney tubular cells we downregulated uPAR expression in kidney proximal tubular epithelial cell line HK-2 by means of cell nucleofection with siRNA. HK-2 cells nucleofected with control Si RNA (SiCo) expressed IL-6 and IL-8 in response to LPS treatment. Similar to the above data, we observed strong downregulation of cell response to LPS in uPARsi cells at mRNA and protein level (Fig. 4A, B).

**Figure 4.**
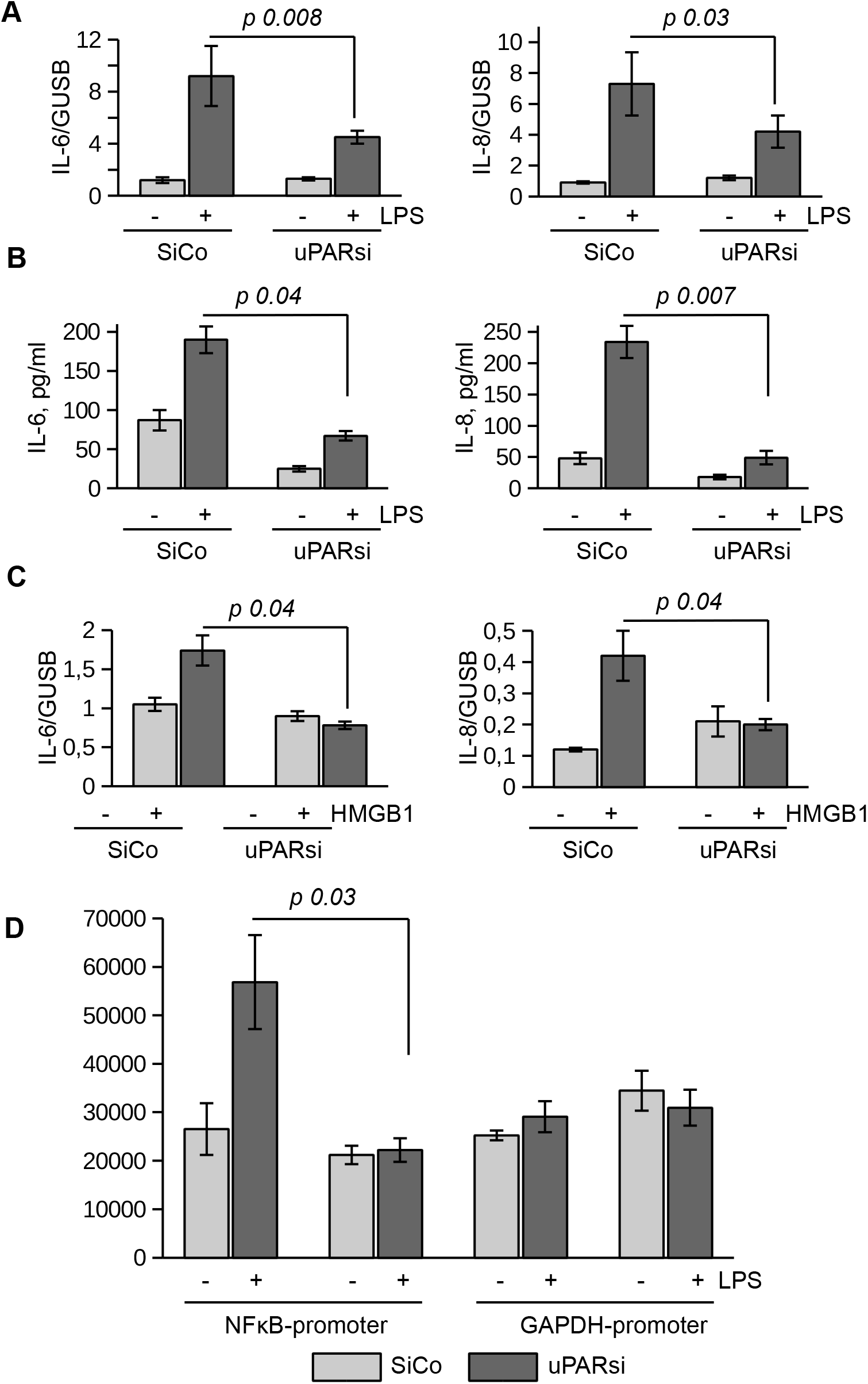
uPAR is essential for the response of kidney proximal tubular epithelial cells to LPS. **A, B.** LPS-induced expression of IL-6 and IL-8 by human renal proximal tubule epithelial cell (HK-2) was assessed by TaqMan RT-PCR (A and ELISA (B). **C.** HMGB1-dependent IL-6 and IL-8 expression by HK-2 cells was assessed by TaqMan RT-PCR. **D.** Human renal epithelial HK-2 cells were lentivirus-infected to express Gaussia luciferase under control of NFκB and GAPDH promoters. Enzyme activity was measured in cell conditioned media 10 hrs after stimulation with LPS.

TLR4 mediates not only signaling induced by PAMPs but is also involved in the recognition of danger associated molecular pattern molecules (DAMPs). One of the important DAMPs is HMGB1 - a DNA binding protein that can be released from damaged cells under stress and activate tubular epithelial cells by interacting with TLR4 in sepsis^24^. In HK-2 cells HMGB1 also induced increased expression of IL-6 and IL-8. Similar to LPS, this response was abrogated in uPARsi cells (Fig. 4C). To assess effects of upAR on LPS-dependent NFκB activation, we infected HK-2 cells with lentivirus to express Gaussia luciferase under control of NFκB-dependent promoter. Gaussia luciferase activity assay showed that LPS-dependent regulation of NFκB-sensitive promoter is dependent on uPAR expression (Fig. 4D) whereas GAPDH promoter is regulated independently on uPAR and LPS.

Together, the above data show that uPAR is involved in mediating LPS-induced effects of TLR4 in myeloid and non-myeloid cells.

### 3. uPAR is a part of TLR4 interactome

Looking for possible mechanisms of uPAR interaction with TLR4 interactome, we found that addition of uPA or blocking uPA/uPAR interaction with antibody had not affected HK-2 cell response to LPS (Supplementary Fig. S2A) suggesting that the observed role of uPAR is independent on its plasminogen activator activity. Downregulation of uPAR expression had also minimal effect on binding of biotin-LPS by these cells as was assessed by the dot blot analysis using biotin-LPS (Fig. 5A).

**Figure 5.**
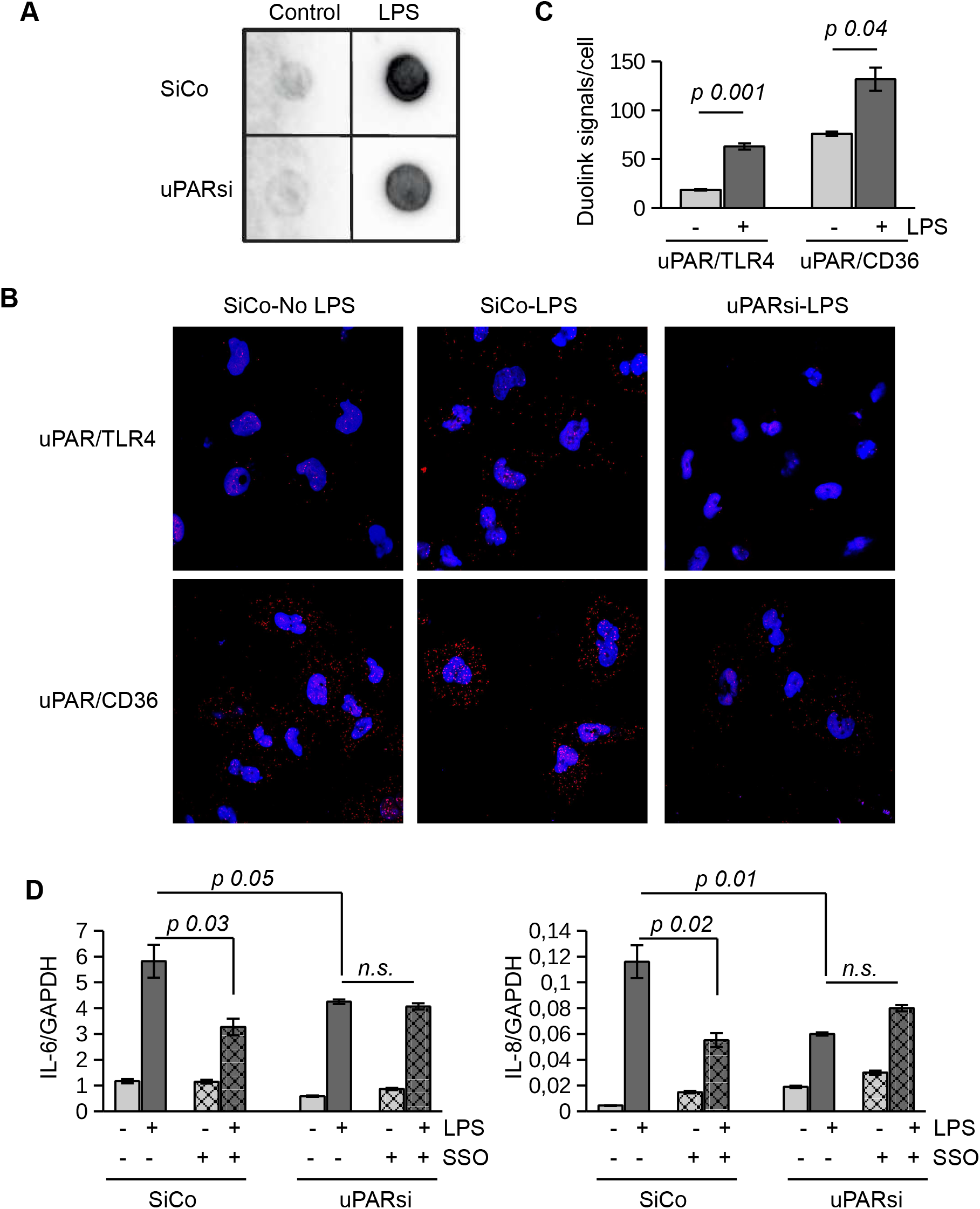
uPAR is a part of TLR4 interactome. **A.** Biotin-LPS binding was assessed in SiCo and uPARsi HK-2 cells as described. **B.** Duolink proximity ligation assay to assess uPAR/TLR4 and uPAR/CD36 interaction was performed on HK-2 cells stimulated with LPS for 15 min as described in Methods. **C.** Duolink images were quantified using Particles analysis tool of ImageJ. **D.** HK-2 cells were stimulated with LPS for 3 hrs after cell pre-treatment with CD36 inhibitor SSO. Expression of IL-6 and IL-8 was assessed by TaqMan RT-PCR.

We performed duolink proximity ligation assay in HK-2 cells to assess the possibility of direct uPAR/TLR4 interaction (Fig. 5B). The number of Duolink signal spots per cell was quantified using ImageJ Analyze particles tool. Relatively weak direct contact observed in control unstimulaated cells was increased in the presence of LPS (Fig 5C). Significantly stronger direct association detected between uPAR and CD36 in unstimulated cells was also further increased by LPS (Fig. 5B, C). Several reports indicated that scavenger receptor CD36 can participate in LPS-induced signaling^18,25^. In our previous work we demonstrated that uPAR cooperates with TLR4 and CD36 to mediate oxLDL signaling in vascular smooth muscle cells. To investigate whether this mechanism can function in LPS signaling, we pre-treated HK-2 cells with CD36 inhibitor SSO prior to LPS stimulaiton. Figure 5D shows that downreguation of uPAR and inhibition of CD36 decreased LPS response in HK-2 cells. However, there was no additive effect of uPARsi and CD36 inhibition suggesting that the receptors are involved in the same signaling mechanism.

### 4. Inflammatory response of uPAR−/− mice is strongly diminished in multimicrobial CLP sepsis model

Immunohistochemical staining showed that uPAR is expressed in vivo in mesothelium of healthy mice whereas the expression of TLR4 was very low. After intraperitoneal (ip) injection of LPS the expession of TLR4 was strongly increased and an association between uPAR and TLR4 could be observed (Fig. 6A) suggesting a possible involvement of uPAR into TLR4 interactome in vivo.

**Figure 6.**
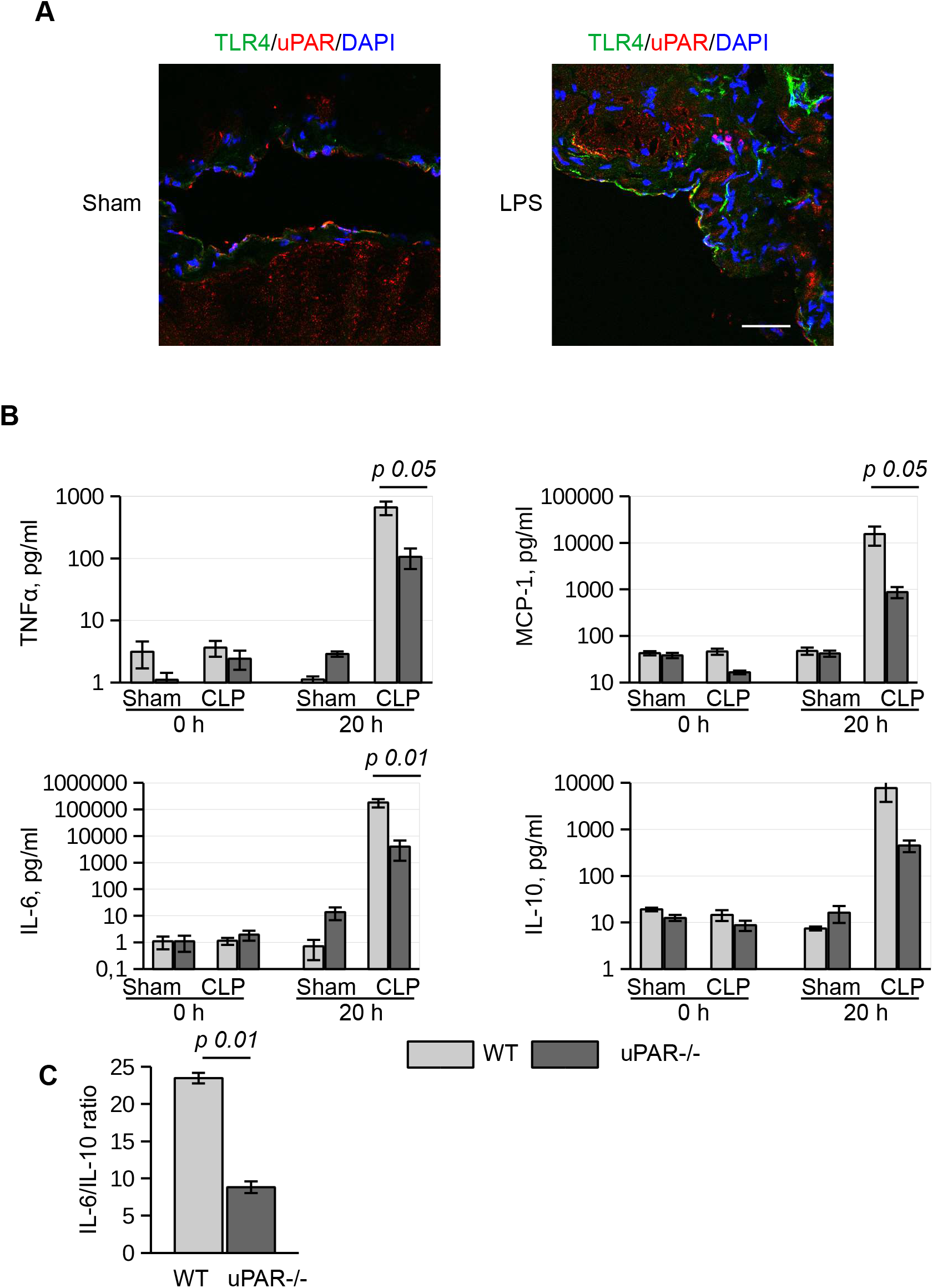
Expression of inflammatory mediators after CLP sepsis is decreased in plasma of uPAR−/− mice. **A.** Peritoneum of sham and LPS-injected WT mice was fixed and stained for uPAR and TLR4. DAPI used as nuclear stain-scale bar 100μm. **B.** Expression of TNFα, MCP-1, IL-6 and IL-1 0 was assessed in mouse blood plasma before and 20 h after CLP surgery using Cytometric Beads Array. C: IL-6/IL-10 ratio in CLP mice 20 hrs after surgery.

The role of uPAR in the inflammatory responses in vivo was further investigated in WT and uPAR−/− mice using the cecal ligation and puncture (CLP) polymicrobial sepsis model. Expression of pro-inflammatory mediators was analyzed in plasma and peritoneal lavage fluid (PLF) 20 hrs after CLP or sham surgery. As expected, CLP-induced peritonitis was associated with a strong local and systemic up-regulation of the pro-inflammatory cytokines IL-6, MCP-1 and TNFα in WT mice. This response was strongly decreased in uPAR−/− mice by 8 and 12 folds for TNFα and MCP-1, respectively in comparison to WT mice (Fig. 6B). IL-6 expression was also low and IL-6/IL10 ratio was also statistically significantly lower in uPAR −/− animals (Fig. 6C). Similar decrease of expression of TNFα, MCP-1, and IL-6 was observed in peritoneal lavage fluid (PLF) performed 20 h after surgery (Fig. 7A).

**Figure 7.**
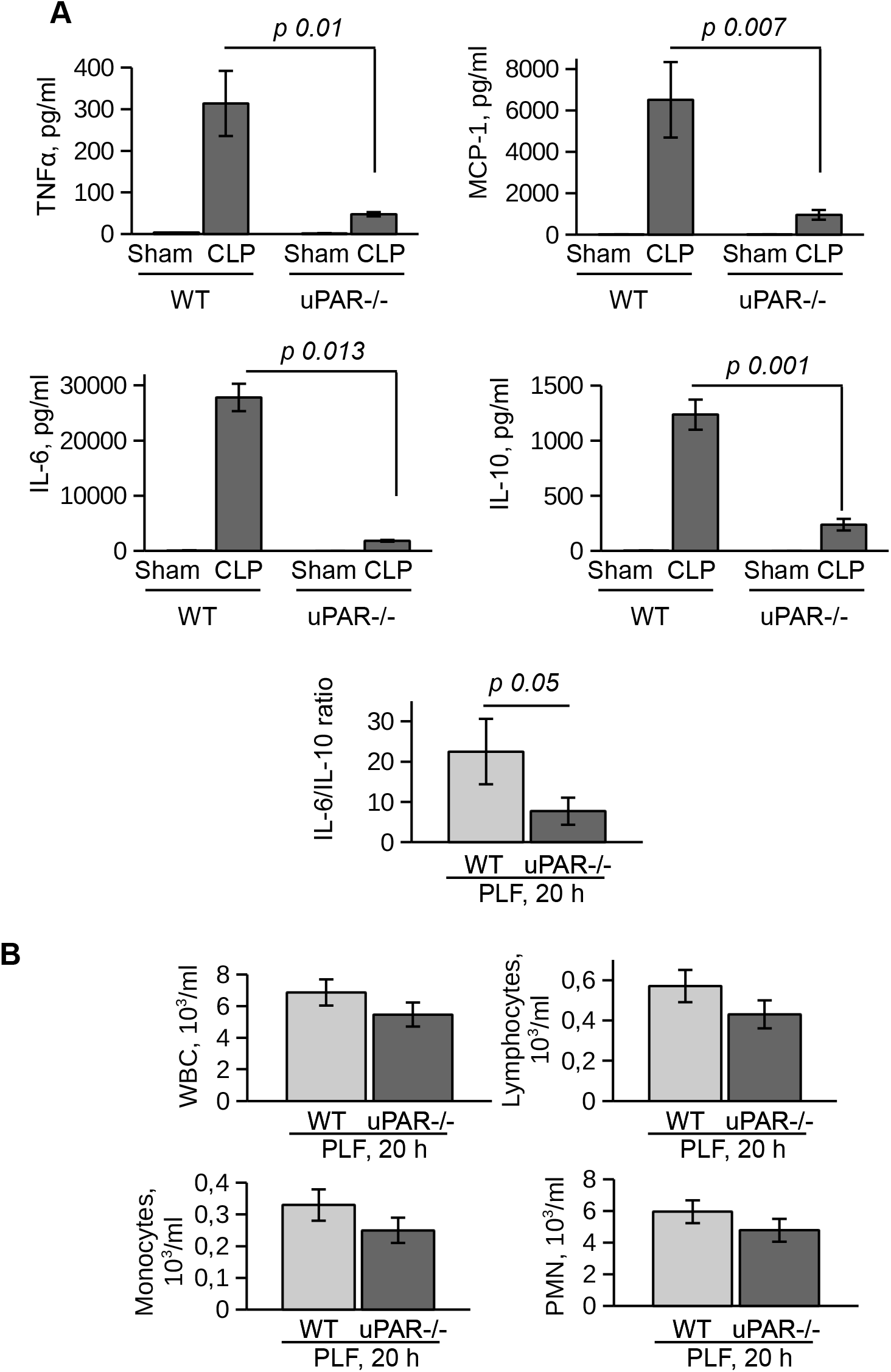
Expression of inflammatory mediators after CLP sepsis is decreased in PLF fluid of uPAR−/− mice. **A.** Expression of TNFα, MCP-1, IL-6 and IL-10 was assessed in PLF 20 h after surgery. **B.** Infiltrating cells in PLF were analyzed 20 h after CLP surgery.

To investigate whether infiltration of inflammatory cells to the peritoneum was impaired in uPAR−/− mice, we analyzed the total number of white blood cells (WBC), as well as quantified the number of lymphocytes, monocytes, and polymorphonuclear leukocytes (PMN) in the blood and PLF 20 h after surgery. The number of WBC decreased in the septic blood in both WT and uPAR −/− mice in a similar way (Supplementary Fig. 3A). Similar to Renckens and colleagues^26^, we have not detected significant deviation in the number of infiltrating inflammatory cells in the peritoneum of uPAR−/− mice in comparison to WT 20 h after CLP surgery (Fig 7B).

Confirming diminished development of inflammatory response, kidney function was significantly improved in uPAR−/− mice. Thus, blood level of creatinine was strongly increased in septic WT mice but remained at the normal level in uPAR−/− mice (Fig. 8A). The level of blood urea nitrogen (BUN) was also significantly diminished in septic uPAR−/− mice compared to WT animals. Liver disfunction was assessed on the basis of enzymatic activities of circulating liver enzymes glutamate oxaloacetate transaminase/aspartate glutaminase (GOT/AST) and glutamate pyruvate transaminase/alanine aminotransferase (GPT/ALT)^27^. Both, GOT and GPT levels were also lower in uPAR−/− septic mice in comparison to WT animals though the differences have not reached statistical significance (Fig. 8B). Plasma level of lactate dehydrogenase (LDH) reflecting the degree of overall tissue damage was also strongly decreased in uPAR−/− mice (Fig. 8C). Basal levels of creatinine, BUN, GOT/AST and GPT/ALT as well as LDH were similar between WT and uPAR−/− mice (baseline in Fig. 8A-C).

**Figure 8.**
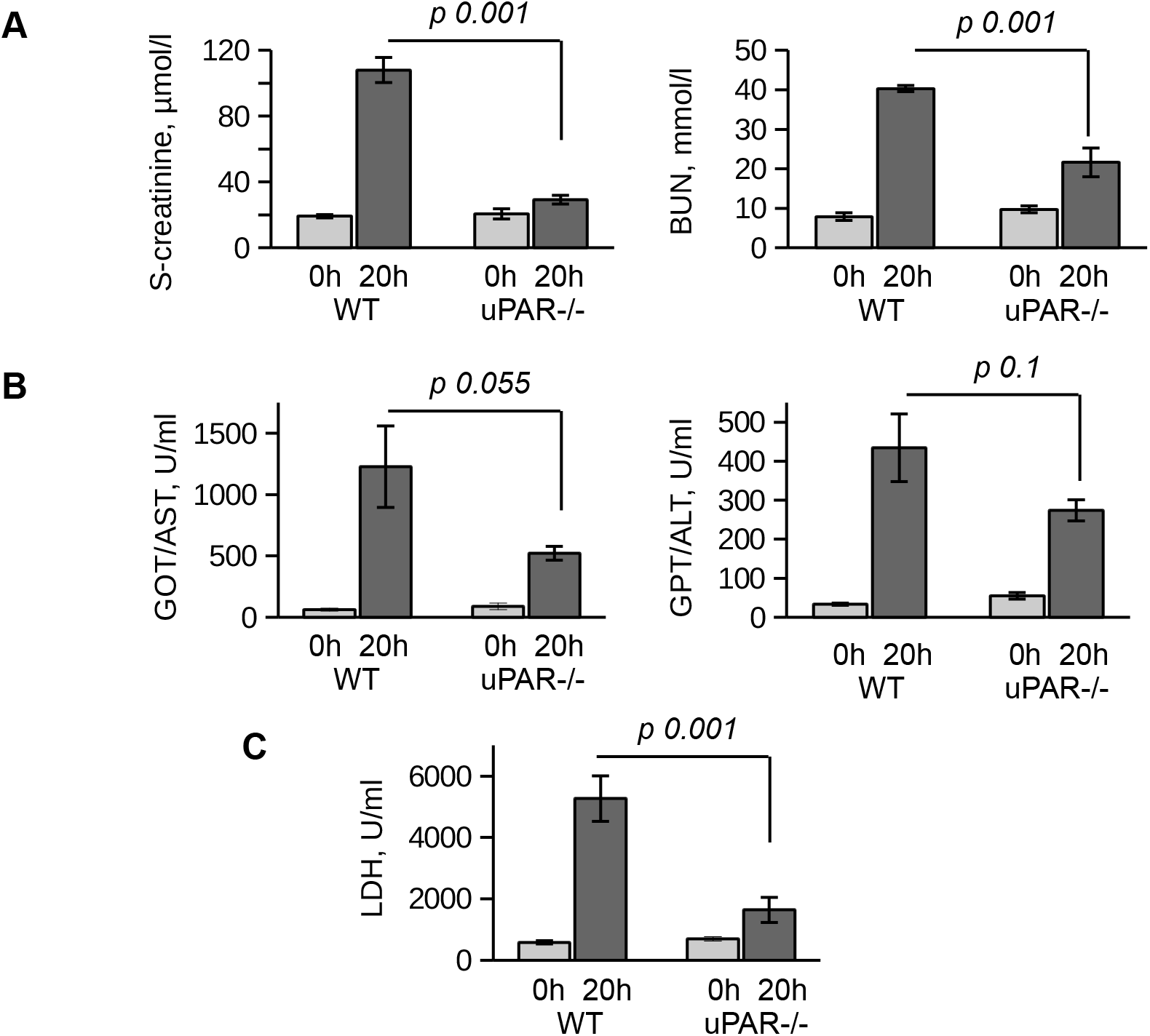
Organ damage in mouse CLP sepsis model is ameliorated in uPAR−/− mice. **A.** Kidney function in WT and uPAR−/− mice at 0 h and 20 hrs after CLP surgery was assessed by analyzing S-creatinine and blood urea nitrogen (BUN). **B.** The function of the liver in CLP mice was assessed by analyzing GOT/AST and GPT/ALT activity in plasma. **C.** and LDH was measured in mouse blood plasma before and 20 hrs after CLP surgery as described in Methods.

## Discussion

In this study we demonstrated that uPAR promotes TLR4-mediated response to LPS in myeloid and non-myeloid cells. Using different approaches, we showed that GPI-uPAR and suPAR can integrate into TLR4 interactome and promote cell signaling leading to the secretion of cytokines and chemokines.

The role of uPAR in immunity is multifaceted and mechanisms are not completely understood. Well-documented is the involvement of uPAR in the migration of inflammatory cells. Migration of granulocytes to the lungs upon pneumococcal pneumonia was impaired in uPAR−/− mice^10^. This was accompanied by increased bacterial load and higher mortality. Similar impairment of neutrophil migration to the lung was also reported upon *Pseudomonas aeruginosa* infection^11^. This was also accompanied by diminished bacterial clearance. Another study showed that during borellia burgdorferi skin infection, the number of spirochetes was increased in uPAR−/− mice^28^. However, in that case infiltration of macrophages was higher in uPAR−/− mice and the effect was attributed to the impaired phagocytosis of bacteria. The mechanisms of uPAR involvement may include regulation of proteolysis on the leading edge of migrating cell^29^chemotaxis and activation of immune cells^30^, due to uPAR interactions with cell surface partners, as integrins and the chemotaxis fMLF-receptors^31^. In addition, uPAR occupation by inactive uPA or its aminoterminal fragment may regulate several activities, including cell adhesion and migration^1^.

Non-proteolytic effects of uPAR on innate immunity were investigated by Liu et al.^13^ during in vitro stimulation of uPAR−/− granulocytes with TLR2 and TLR4 ligands. They showed that uPAR is essential for cell response to TLR2 ligand. The mRNA expression was not decreased, however, release of IL-6 and TNFa was diminished in uPAR−/− cells. The authors also stimulated uPAR−/− granulocytes with TLR4 ligand LPS and observed no changes in mRNA expression of IL-6 and TNFa after 24 hrs of stimulation. On the contrary, in our experiments we demonstrated diminished mRNA expression of LPS-induced inflammatory mediators in different cell types after downregulation of uPAR. The discrepancy with the data by Liu et al. can be explained by different stimulation conditions. In our experiments stimulation with LPS for 3 hrs was found optimal to assess changes of mRNA expression. Changes of protein expression were pronounced 24 hrs after treatment, whereas the changes of RNA expression were no longer visible at that time point. Similar to Liu and colleagues, in our experiments TLR1 and TLR1/2 ligands PAM3CSK4 and lipoteichoic acid (LTA) also induced less inflammatory response in uPARsi cells (data not shown).

Our data showed that uPAR interferes with signaling of TLR4 to different PAMP and DAMP molecules. We also found uPAR to be a part of TLR4 interactome. Interestingly, suPAR also interacted with TLR4 and promoted LPS signaling in uPAR−/− cells. These data suggest that regulation of LPS response of TLR4 by (s)uPAR depends on the availability of membrane-bound and soluble uPAR. LPS signaling of TLR4 is very complex. In addition to CD14 and MD2 co-receptors, recent data demonstrated that TLR4 can recruit further membrane receptors such as TLR2, CD36, integrin CD11b, heat shock proteins and others^32^. Looking for possible mechanisms of uPAR effects, we showed that these effects were not dependent on uPA/uPAR interaction. Rather, TLR4 or uPAR interaction with common co-receptors was affected. Recent report showed that LPS-dependent signaling and expression of inflammatory mediators was decreased after silencing CD36 in epithelial cells^25^. Accordingly, we showed that inhibition of CD36 decreases LPS response in SiCo but not in uPARsi HK-2 cells, suggesting that uPAR and CD36 are involved in the same molecular mechanism. So, it is possible that uPAR mediates LPS-dependent TLR4/CD36 cross-talk in a similar fashion as we have previously demonstrated for oxLDL signaling^17^. This hypothesis is strengthened by our observation that expression of uPAR did not promote LPS response in HEK-BlueTLR4 reporter cell line (Invivogen) – HEK 293 cells that stably express TLR4, CD14 and MD-2. It is recognized that transcriptome of HEK 293 is specific and the cells do not express a variety of scavenger receptors and PRRs. This data suggest that fine mechanisms of (s)uPAR interference is probably cell type dependent, depend on the ligand nature, and can be fine-tuned by the availability of various co-receptors, and further factors.

In vivo in CLP mouse polymicrobial sepsis model we found strongly decreased level of TNFa, MCP-1, and IL-6 in PLF and in blood plasma of uPAR−/− mice. Interestingly, recruitment of innate immunity cells to the peritoneum was similar between uPAR−/− and WT mice 20 hrs after surgery. This data is in agreement with the report by Renckens et al.^26^, who showed that LPS-dependent migration after ip LPS injection was impaired in uPAR−/− mice, whereas the effects were compensated upon sepsis induction by the injection of living E-coli. Importantly, we also found that plasma content of LDH indicating overall tissue damage in sepsis was strongly decreased in uPAR−/− mice. Kidney function was also improved as was assessed by the plasma content of creatinine and BUN. We also observed a trend towards improvement of the liver function though these differences have not reached statistical significance. It should be kept in mind that CLP is a very severe sepsis model where inflammation is induced by combination of gram-positive and gram-negative bacteria that can be recognized by many pattern recognition receptors. An important role of TLR4 in this process was confirmed by the observation of Chen et al. who showed that TLR4−/− mice demonstrated improved survival and decreased level of cytokines after CLP^33^. Knockdown of CD14 that functions not only with TLR4 but also with other TLRs had even stronger protective effect in mouse CLP model^34^. Our data show that uPAR that can interfere with PRRs signaling and thus promote immune response. This mechanism represents a potentially important target in sepsis therapy. Further research is needed to identify uPAR interaction partner-PRR and develop a strategy to target this interaction.

## Methods

### Materials

Unconjugated and Alexa 647-conjugated mouse uPAR antibody were from R&D systems (MAB531 and FAB531R, respectively); TLR4 antibody (MAB27591) was from SrnD Systems; LPS (L2887) was from Sigma, Biotin-LPS and PAM3CSK4 were from Invivogen; Soluble mouse uPAR was from CinoBiologicals. Human IL-6 and IL-8 ELISAs were from Thermofisher Scientific. Mouse inflammation CBA kit was from BD Biosciences. Mouse and human uPAR siRNA and non-sence siRNA control duplexes were from Santa Cruz Biotechnology. *RT-PCR*. RNA was isolated using RNAEasy kit from Quiagen. TaqMan RT-PCR was performed using TaqMan Master Mix and Light Cycler96 (Roche). Oligonucleotides are listed in supplementary Table 1.

### Cell culture, transfection, and luciferase assay

Immortalized mouse peritoneal mesothelial cell line was generated in our lab by limited dilution cultures of primary cells obtained from omentum tissue of Immorto mice harboring the tsSV40T gene as previously described^35^. The cells were propagated in RPMI 1640 cell culture medium containing 1% penicillin–streptomycin, 10% fetal calf serum, 1% insulin/transferrin/selenium A (all from Life Technologies, Carlsbad, CA), 0.4 μg/ml hydrocortisone (Sigma-Aldrich), and 10 U/ml recombinant mouse interferon gamma (Cell Sciences, Canton, MA) at 33 °C (permissive conditions). The cell lines were identified by the typical cobblestone morphology of confluent monolayers and by positive staining for E-cadherin, ZO-1, α-SMA, and pan-cytokeratin after 3-day culture at 37 °C without interferon gamma (non-permissive conditions). Primary preitoneal mouse macrophages were isolated from uPAR−/− mice as described previously^36^.

Raw 264.7 mouse macrophage cell line was from ATCC and cultivated as recommended by the supplier. HK-2 human kidney proximal tubule epithelial cells were from ATCC and cultivated as recommended by the supplier in Keratinocyte Serum Free Medium containing 0.05 mg/ml bovine pituitary extract and 5 ng/ml EGF.

Mesothelial cells were transfected using PolyPlus transfection reagents accordingly to the manufacturer instructions. HK-2 cells were nucleofected using Mirus nucleofection solution and T20 program of nucleofector (Lonza).

Construction of vector for Gaussia luciferase expression under control of NFκB promoter was described elsewhere^17^. Activity was measured using GeneCopoeia kits and Tecan Genios multiplate reader.

### Ex vivo blood stimulation

WT and uPAR−/− mouse hole blood was collected in EDTA tube. Stimulation was performed with 50 ng/ml LPS for 3 hrs at 37°C. Then, the blood was centrifuged at 1500 g for 10 min, and plasma was used for cytokines measurement using Cytokine Beads Array.

### Biotin-LPS binding, pull down and western blotting

To analyze LPS binding, 1 μg/ml Biotin-LPS was added to the cells. After 30 min of incubation, cells were washed and lysed. For Dot Blot analysis, 10μg of cell lysate protein was applied to nitrocellulose membrane. Membrane was allowed to dry, blocked in 3% BSA, and incubate with Streptavidine-HRP for 1 h at room temperature. After washing, membrane was developed using Versa Doc Gel Documentation system (BioRad) and QuantityOne software. For pull-down assay, cell lysate was incubated with streptavidine magnetic beads. Beads were then washed, SDS electrophoresis and western blotting have been performed.

### Immunocytochemistry and immunohistochemistry

Cells were grown on coverslips and stimulated as indicated. The cells were fixed and processed for immunostaining as we have previously described. Staining with antibodies was performed for 1 h at room temperature. DAPI was applied for nuclear staining. Duolink proximity assay kit was purchased from Sigma and used as recommended by the supplier. Leica TCS-SP2 AOBS confocal microscope (Leica Microsystems). All the images were taken with oil-immersed x40 objective, NA 1.25 and x63 objective, NA 1.4.

### Animal experiments

All procedures were performed in accordance with international guidelines on animal experimentation and approved by the local committee for care and use of laboratory animals (Lower Saxony Office for Consumer Protection and Food Safety). Experiments were performed as previously described^37^. Briefly, wild type C57BL/6J and uPAR−/− B6.129P2-*Plaur*^*tm1Jld*^/J male mice (20 to 25 g) obtained from Charles River Laboratories (Sulzfeld, Germany) were anesthetized with isofluorane (induction of 3%, maintenance of 1.5%, and oxygen flow of 3 L/minute). A 1-cm ventral midline abdominal incision was made and the cecum was ligated with 4-0 silk sutures just distal to the ileocecal valve and punctured through with a 24-gauge needle. 1- to 2-mm droplet of fecal material was expelled. The incision was closed using 4-0 surgical sutures. Mice were fluid-resuscitated with 500μL prewarmed normal saline intraperitoneally immediately after the procedure. Sham animals underwent the same procedure except for CLP. Mice were anesthetized for blood sampling with isofluorane and subsequently sacrificed at 18 hours after CLP or sham surgery (n = 10 per group). Serum level of creatinine and urea and the activities of GOT/AST and GPT/ALT were measured by an automated method and an Olympus AU 400 analyzer (Beckman Coulter Inc., Krefeld, Germany). Serum levels of the tumor necrosis factor-alpha (TNF-a), interleukin-6 (IL-6), and macrophage chemoattractant protein-1 (MCP-1) were quantified by bead-based flow cytometry assay (CBA Kit; BD Biosciences, Heidelberg, Germany) in accordance with the instructions of the manufacturer.

### Statistics

Data are shown as mean ± SEM throughout this study. All experiments were independently repeated at least three times. Student’s T test test was applied for comparing two different groups of data. Multiple comparisons were analyzed by using the one-way analysis of variance with the Tukey as a *post hoc* test. GraphPad Prism version 5.02 (GraphPad Prism Software Inc., San Diego, CA, USA) was used for data analysis.

