## Supplementary figs for "Urokinase receptor associates with TLR4 interactome to promote LPS response"

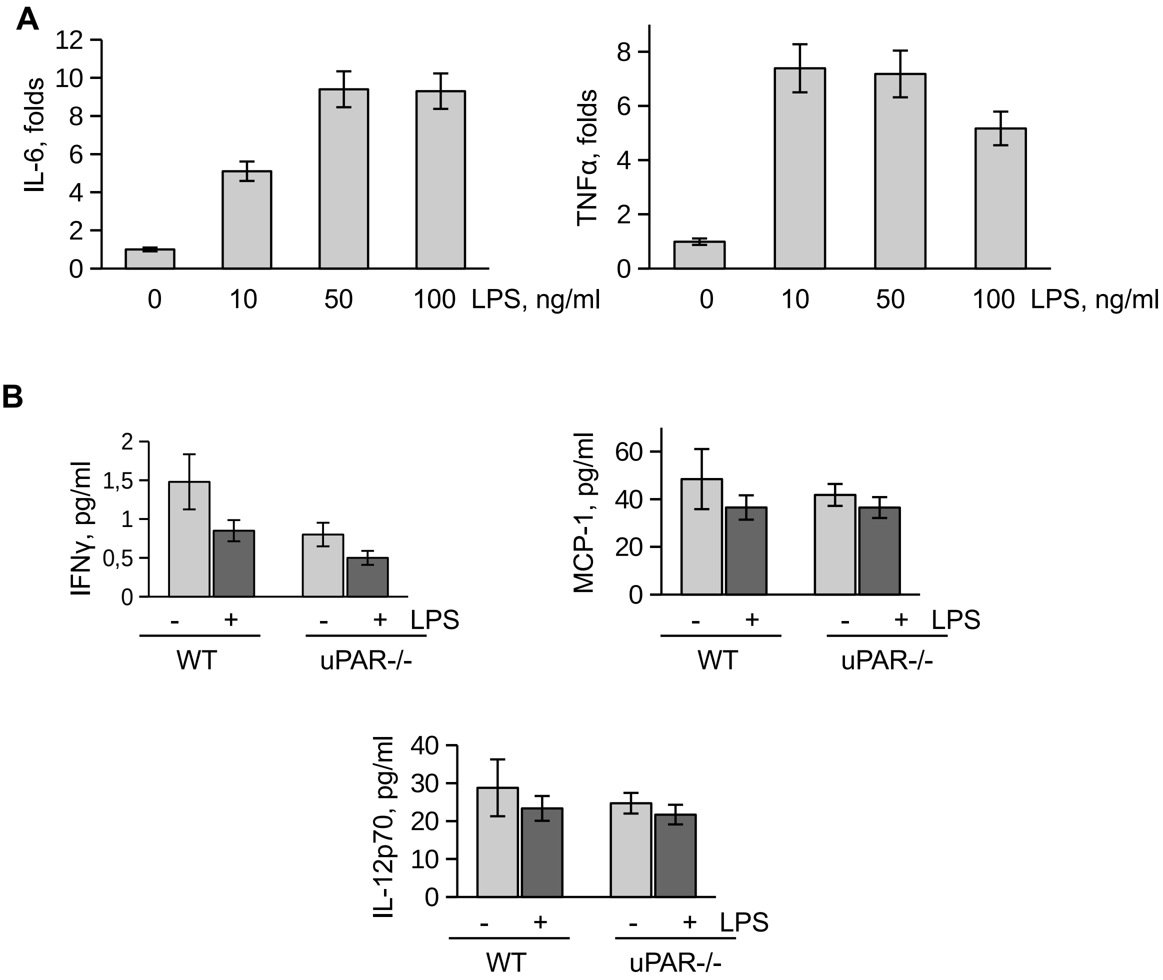


**Supplementary figure S1.** A. Optimization of the ex vivo mouse blood stimulation with LPS. Mouse blood was stimulated ex vivo with 10, 50, 100 ng/ml LPS for 2 h.IL-6 and TNFα was measured using CBA kit. B. IFNγ, MCP-1, and IL-12p70 measured after ex vivo LPS stimulation of WT and uPAR-/- blood.

A

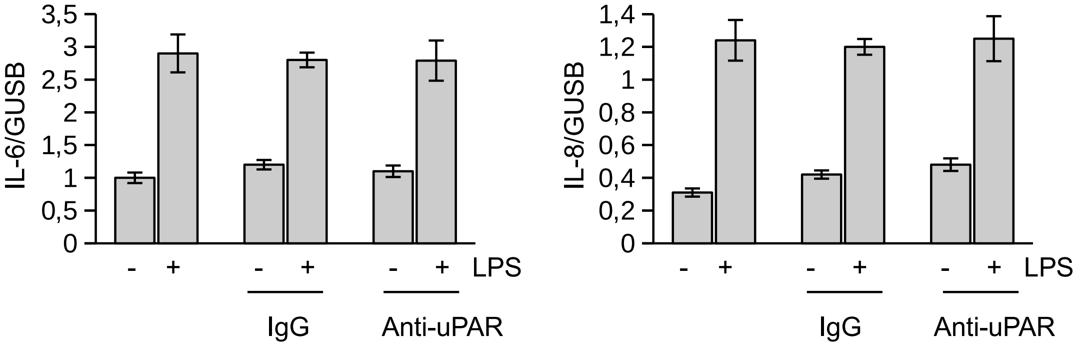


**Supplementary figure S2.** A. HK-2 were pre-treated with anti-uPAR antibody or IgG before LPS stimulation. Expression of IL-6 and IL-8 was assessed by RT-PCR.

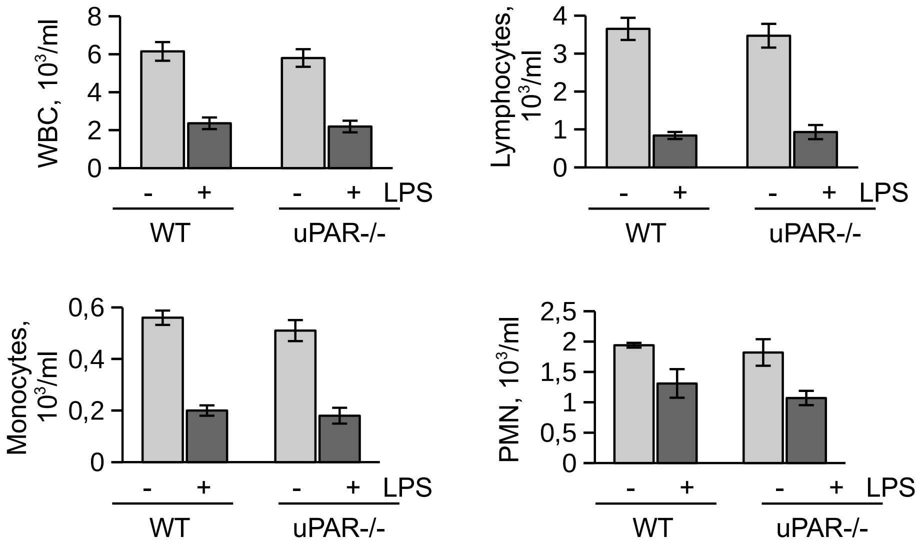


**Supplementary figure S3.** Blood cells count in WT and uPAR-/- 20 hrs after CLP surgery.
